# Quantifying synthetic bacterial community composition with flow cytometry: efficacy in mock communities and challenges in co-cultures

**DOI:** 10.1101/2024.07.26.605308

**Authors:** Fabian Mermans, Ioanna Chatzigiannidou, Wim Teughels, Nico Boon

## Abstract

Determination of bacterial community composition in synthetic communities is critical for understanding microbial systems. The community composition is typically determined through bacterial plating or through PCR-based methods which can be labor-intensive, expensive or prone to bias. Simultaneously, flow cytometry has been suggested as a cheap and fast alternative. However, since the technique captures the phenotypic state of bacterial cells, accurate determination of community composition could be affected when bacteria are co-cultured. We investigated the performance of flow cytometry for quantifying oral synthetic communities and compared it to the performance of strain specific qPCR and 16S rRNA gene amplicon sequencing. Therefore, axenic cultures, mock communities and co-cultures of oral bacteria were prepared. Random forest classifiers trained on flow cytometry data of axenic cultures were used to determine the composition of the synthetic communities, as well as strain specific qPCR and 16S rRNA gene amplicon sequencing. Flow cytometry was shown to have a lower average root mean squared error and outperformed the PCR-based methods in even mock communities (flow cytometry: 0.11 ± 0.04; qPCR: 0.26 ± 0.09; amplicon sequencing: 0.15 ± 0.01). When bacteria were co-cultured, neither flow cytometry, strain specific qPCR and 16S rRNA gene amplicon sequencing resulted in similar community composition. Performance of flow cytometry was decreased compared to mock communities due to changing phenotypes. Finally, discrepancies between flow cytometry and strain specific qPCR were found. These findings highlight the challenges ahead for quantifying community composition in co-cultures by flow cytometry.

**Importance:** Quantification of bacterial composition in synthetic communities is crucial for understanding and steering microbial interactions. Traditional approaches like plating, strain specific qPCR and amplicon sequencing are often labor-intensive and expensive and limit high-throughput experiments. Recently, flow cytometry has been suggested as a swift and cheap alternative for quantifying communities and has been successfully demonstrated on simple bacterial mock communities. However, since flow cytometry measures the phenotypic state of cells, measurements can be affected by differing phenotypes. Especially changing phenotypes resulting from co-culturing bacteria can have a profound effect on the applicability of the technique in this context. This research illustrates the feasibility and challenges of flow cytometry for the determination of community structure in synthetic mock communities and co-cultures.

## Introduction

Microbial communities play central roles in environmental and industrial processes, as well as in animal and human health. Elucidating their functionality and structure is key to understanding and steering these processes (1–4). Specifically for the oral microbiome, shifts in the microbial community influence the host health status and progression of disease, highlighting a need to study these phenomena (4–7). To better apprehend these interactions, *in vitro* models have proven useful, allowing controlled manipulation of the system. Many of these models use low diverse synthetic microbial communities, consisting of a few different bacteria. This allows for easier mechanistic research on the interactions of the different community members, their environment and the host (8–16).

Analysis of the community composition in these synthetic microbial systems is typically done by plate counting or PCR-based methods (17–21). Plate counting suffers from bias caused by an adopted viable but nonculturable (VBNC) state of lab cultures. This leads to underestimation of the population density, often referred to as the great plate count anomaly (22, 23). Additionally, the technique requires that each member of the community can be differentiated from the other members by using selective media or by assessment of morphological differences and is labor-intensive (24, 25). PCR-based methods, with most notably quantitative PCR (qPCR) and 16S rRNA gene amplicon sequencing, have shortcomings as well. Communities consisting of different taxa are prone to bias because of taxon-dependent nucleic acid extraction efficiencies, primer selectivity, and varying amplification efficiency (26–30). Strain-specific qPCR requires adequate probe and primer design, which is often a time-consuming process. Additionally, the number of strains that can be quantified by multiplex qPCR in a single run is restricted due to the limited number of fluorophores that can be detected by the instrument, leading to increased analysis time (31, 32). With 16S rRNA gene amplicon sequencing, the limited length of targeted fragments can be an issue for taxonomic resolution when using closely related organisms, some form of normalization is needed to account for differing sequencing depths, and direct absolute quantification is not possible (33–35). Nevertheless, this last issue can be circumvented by combining the data with other techniques, such as flow cytometry (36).

Recently, flow cytometry has been proposed to identify bacterial strains and quantify microbial community composition (37–40). Flow cytometry is independent of subsequent culturing, does not require a nucleic acid extraction step and provides optical multiparameter data on a single cell level for the entire community in a matter of minutes (41, 42). It is compatible with a wide variety of staining techniques, allowing monitoring of physiological activity as well as specifically target biomolecules (43–46). Application and development of the technique keeps progressing, leading to a growing field of flow cytometry bioinformatics (37, 47–51). Moreover, the technique is cheap, where a sample can be analyzed for less than 1 euro. However, flow cytometry measures the phenotypic state of cells, which can lead to different fingerprints for the same bacterial strain when cultured under different conditions or when sampled in different growth phases (38, 52–54). This could prove a problem when trying to determine the composition of bacterial communities, especially when bacteria are co-cultured. Previous studies testing the feasibility of flow cytometry for determination of microbial community structure showed that it is possible to determine the relative community composition of *in silico* and *in vitro* mock communities for simple microbial consortia but were unable to assess its capabilities towards co-cultures (37, 40, 55).

In this study we investigated whether flow cytometry can be used for determining the relative and absolute community composition of oral synthetic communities, both mock communities and co-cultures. For this purpose, we prepared axenic cultures, *in silico* mock communities, *in vitro* mock communities and co-cultures of oral bacteria. Flow cytometry data of axenic cultures was used to train random forest classifiers, which in turn were used to predict the composition of the *in silico* and *in vitro* mock communities and co-cultures. Concurrently, DNA extracts from the *in vitro* mock communities and co-cultures were used to determine to composition by strain specific qPCR and 16S rRNA gene amplicon sequencing.

## Materials and Methods

A general overview of the workflow is provided in Figure 1.

**Figure 1.**
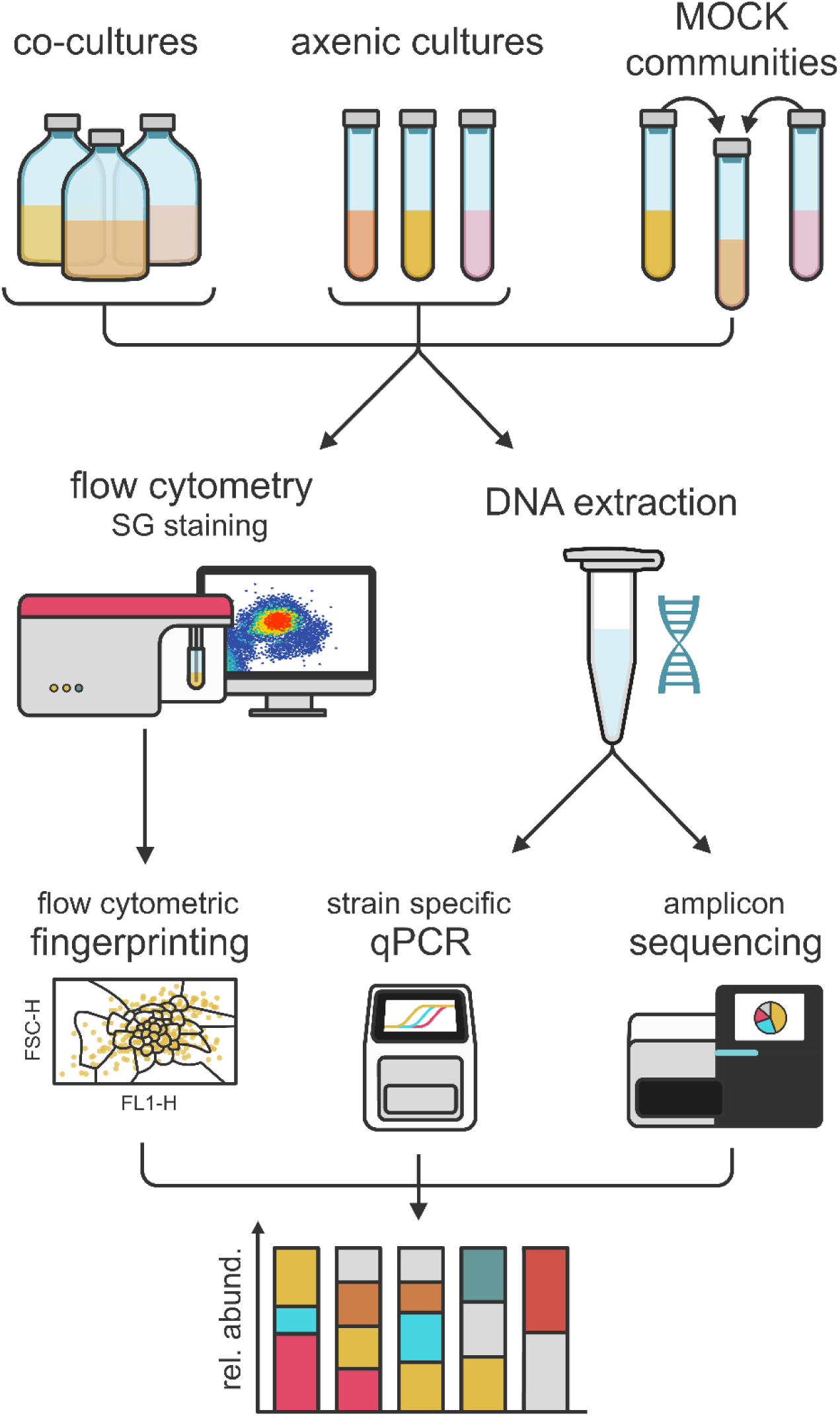
Overview of the workflow. Axenic cultures, mock communities and co-cultures of oral bacteria were prepared. In turn, the community composition was determined through flow cytometry, strain specific qPCR and 16S rRNA gene amplicon sequencing.

### Strains and growth conditions

*Aggregatibacter actinomycetemcomitans* ATCC 43718, *Actinomyces naeslundii* ATCC 51655, *Actinomyces viscosus* ATCC 15987, *Fusobacterium nucleatum* ATCC 10953, *Porphyromonas gingivalis* ATCC 33277, *Prevotella intermedia* ATCC 25611 and *Veillonella parvula* DSM 2007 were maintained on blood agar base No. 2 (Oxoid, Hampshire, UK) supplemented with menadione (1 mg/L) (Sigma-Aldrich, Diegem, Belgium), hemin (5 mg/L) (Sigma-Aldrich, Diegem, Belgium), and 5% sterile defibrinated horse blood (Oxoid, Hampshire, UK) under anaerobic conditions (90% N_2_, 10% CO_2_) at 37°C. All strains maintained on blood agar as well as frozen stocks of *Streptococcus gordonii* ATCC 49818, *Streptococcus mitis* ATCC 49456, *Streptococcus mutans* ATCC 25175, *Streptococcus oralis* ATCC 35037, *Streptococcus salivarius* TOVE-R, *Streptococcus sanguinis* LMG 14657 and *Streptococcus sobrinus* ATCC 33478 were incubated in triplicate in Brain Heart Infusion broth 2 (BHI-2) (56) under anaerobic conditions (90% N_2_, 10% CO_2_) at 37°C for 24h. All bacterial strains were grown in triplicate. After incubation, cell concentrations were determined by flow cytometry and a sample for DNA extraction was taken. BHI-2 was prepared using Brain Heart Infusion (Karl Roth, Karlsruhe, Germany) supplemented with mucin from porcine stomach type III (2.5 g/L) (Sigma-Aldrich, Diegem, Belgium), yeast extract (1 g/L) (Oxoid, Hampshire, UK), cysteine (0.1 g/L) (Merck, Darmstadt, Germany), sodium bicarbonate (2 g/L) (Sigma-Aldrich, Diegem, Belgium), glutamic acid (2.5 g/L) (Merck, Darmstadt, Germany), hemin (5 mg/L) (Sigma-Aldrich, Diegem, Belgium), and menadione (1 mg/L) (Sigma-Aldrich, Diegem, Belgium).

### Mock communities

Mock communities of different bacteria were prepared by mixing the different axenic cultures. *Mock 1-7* were prepared so that the microbial community would be even, *Mock 8-9* were prepared as uneven communities (Figure 2; Supplementary 1). Axenic cultures were vortexed for 5 seconds prior to mixing and the mock communities were homogenized by vortexing for 20 seconds post mixing. Immediately after preparing the mock communities, samples were measured by flow cytometry and samples for DNA extraction were taken.

**Figure 2.**
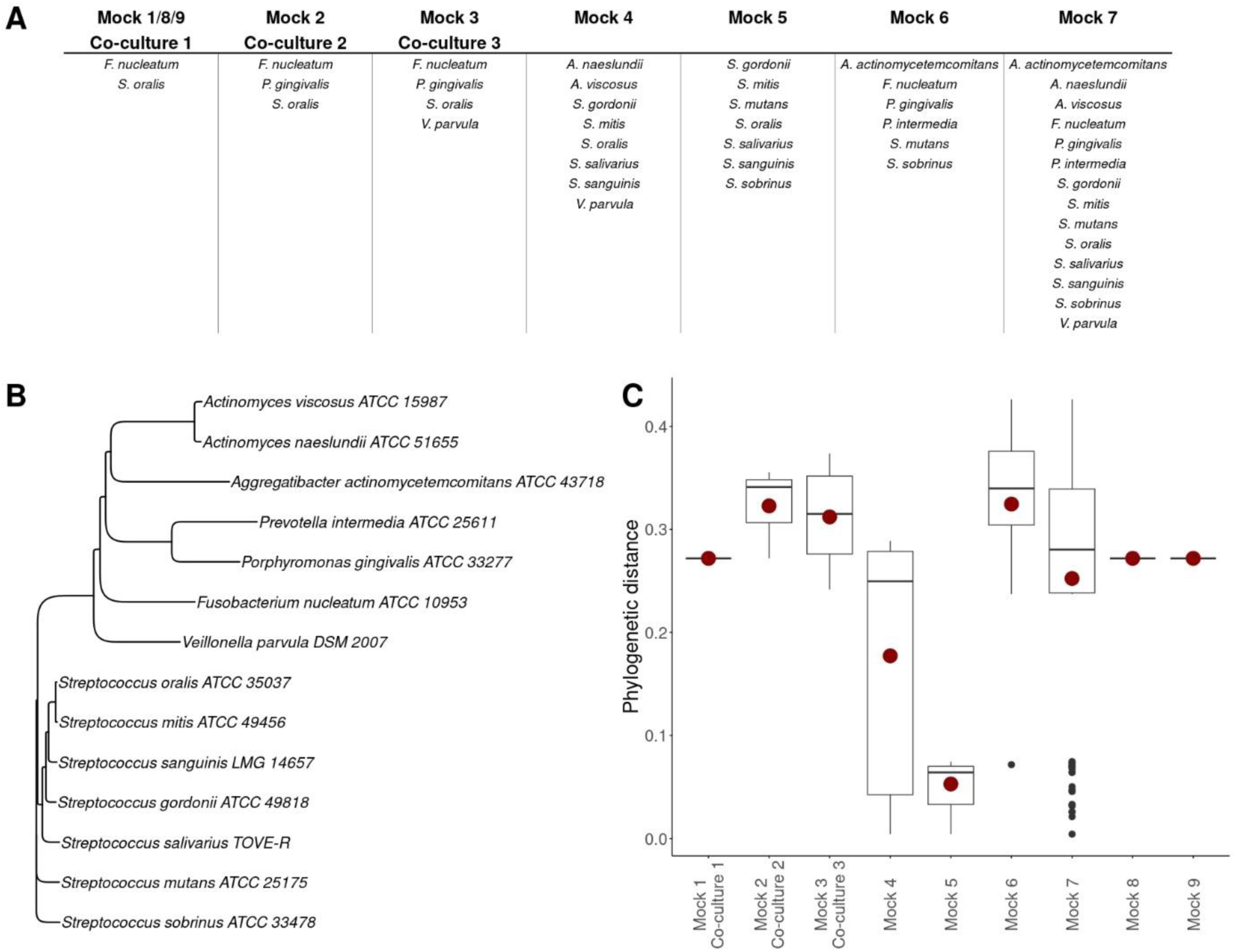
Information on the different mock communities and co-cultures. (A) Community membership of each bacterial strain to the mock communities and co-cultures. (B) Phylogenetic tree of all used bacterial strains. The tree was constructed using the neighbor joining clustering method and based on partial sequences of the 16S rRNA gene obtained through Sanger sequencing. (C) Phylogenetic distance between community members of each synthetic community. The red dot shows the mean phylogenetic distance of each community.

### Co-cultures

24-hour cultures of *S. oralis*, *F. nucleatum*, *P. gingivalis* and *V. parvula* in BHI-2 were inoculated in equal cell concentrations, as determined by flow cytometry, in anaerobic penicillin bottles (90% N_2_, 10% CO_2_) with fresh BHI-2 in triplicate (Figure 2). Consecutively, the mixtures were incubated at 37°C for 48h. After 24h and 48h of incubation, co-cultures were measured by flow cytometry and samples for DNA extraction were taken.

### Flow cytometry

#### Sample preparation

Axenic cultures, mock communities and co-cultures were vortexed for 5 seconds and consecutively diluted 1000-fold in sterile PBS (PBS tablet, Sigma-Aldrich, Steinheim, Germany). Samples then were stained with 1% v/v SYBR® Green I (SG) stock solution and incubated in the dark at 37°C for 20 minutes. The SG stock solution was prepared by diluting SYBR® Green I (10000x concentrate in DMSO) (Invitrogen, Eugene, USA) 100-fold in 0.2 µm filtered DMSO (Merck, Darmstadt, Germany).

#### Sample measurement

Stained samples were measured using an Attune NxT (Invitrogen, Carlsbad, USA) flow cytometer equipped with a blue (488 nm) and red (638 nm) laser. Performance of the instrument was checked using Attune Performance tracking beads (Invitrogen, Eugene, USA). Only the blue laser was used for excitation of the stain. A 530/30 nm band-pass filter was used for the detection of green fluorescence (BL1) and a 695/40 nm band-pass filter was used for the detection of red fluorescence (BL3). A 488/10 band-pass filter was used for the detection of forward scatter (FSC) and side scatter (SSC). The flow rate was set to 100 µL/min and stop conditions were set to 85 µL of sample analyzed. The threshold (set on green fluorescence, BL1) and PMT-voltages were determined by using control samples: an axenic culture, sterile BHI-2, and sterile PBS.

### DNA extraction, qPCR and sequencing

DNA extraction was performed as previously described by De Paepe et al. (57). Samples were centrifuged at 20238 x g for 10 minutes and supernatant was discarded. The remaining pellet was mixed with 1 mL of lysis buffer, containing 100 mM Tris pH 8 (Carl Roth, Karlsruhe, Germany), 100 mM EDTA pH 8 (Carl Roth, Karlsruhe, Germany), 100 mM NaCl (Carl Roth, Karlsruhe, Germany), 1% polyvinylpyrrolidone (PVP40) (Sigma-Aldrich, Steinheim, Germany) and 2% sodium dodecyl sulphate (SDS) (Carl Roth, Karlsruhe, Germany). 400 mg of 0.1 mm glass beads (Omni International, Kennesaw, USA) were added to the samples after which they were disrupted in the PowerLyzer (Qiagen, Venlo, Netherlands) at 2000 rpm for 5 minutes. Samples then were centrifuged at 20238 x g for 5 minutes and supernatant was added to a new tube containing 500 µL of phenol:chloroform:isoamilic alcohol 25:24:1 pH 7 (Thermo Fisher Scientific, Carlsbad, USA). Samples were mixed and centrifuged at 20238 x g for 1 minute. Consecutively, 450 µL of the upper phase was added to a new tube containing 700 µL chloroform (Merck, Darmstadt, Germany). After mixing and centrifuging at 20238 x g for 1 minute, 450 µL of the upper phase was transferred to a new tube containing 500 µL of cold isopropanol (VWR, Fontenay-sous-Bois, France) and 45 µL of 3M sodium acetate (Carl Roth, Karlsruhe, Germany). Samples were mixed and stored at -20 °C for one hour after which they were centrifuged at 20238 x g at 4 °C for 30 minutes. Supernatant was removed and the DNA pellet was air dried prior to dissolving in 1X TE buffer (Rockland, Limerick, USA).

Each bacterial strain was quantified by qPCR in a Taqman 5’ nuclease assay using specific primers and probes (Supplementary 2) (58, 59). qPCR assays were performed using a StepOnePlus real-time PCR system (Applied Biosystems, Foster City, California, USA). Reactions were executed in a volume of 25 µL consisting of 12.5 µL Takyon ROX Probe 2X MasterMix dTTP blue (Eurogentec, Seraing, Belgium), 5 µL DNA template, 1 µL (10 µM stock) of each primer, 1 µL of probe, and 4.5 µL nuclease-free water (Serva, Heidelberg, Germany). Amplifications were run as follows: initial denaturation at 50 °C for 2 minutes and at 95 °C for 10 minutes, followed by 40 cycles of 15 seconds denaturation at 95 °C and combined anneal/extension at 60 °C for 1 minute. A plasmid standard curve was used to perform quantification. All qPCR analyses were done in triplicate. 10 µL genomic DNA extract was send to LGC genomics GmbH (Berlin, Germany) for library preparation and sequencing on an Illumina MiSeq platform with v3 chemistry with the primers 341F (5’ – CCT ACG GGN GGC WGC AG – 3’) and 785Rmod (5’ – GAC TAC HVG GGT ATC TAA KCC – 3’) (60).

### Data analysis

#### Analysis of flow cytometry data

Flow cytometry data were imported in R (version 4.2.0) using the flowCore package (version 2.8.0) (61). Data were transformed using the arcsine hyperbolic function (50), and gated manually on the primary fluorescent channels (BL1 and BL3) to remove background (Supplementary 3). Cell concentrations were determined based on the number of events in the cell gate and the volumetric measurements of the flow cytometer. Singlets were gated manually on side scatter (SSC) by assessing the relationship between peak height (SSC-H) and peak area (SSC-A) (Supplementary 3). Data of replicates of the axenic cultures were pooled to train random forest classifiers. Random forest classifiers were constructed using the ‘*RandomF_FCS’* function of the Phenoflow package (version 1.1.2) (50). The parameters upon which the models were build were FSC-H, FSC-A, SSC-H, SSC-A, BL1-H, BL1-A, BL3-H, and BL3-A because these are expected to contain the most information (39). 75% of the data was used for training the models and the remaining 25% of the data was used for testing. Accuracy was used as performance metric to optimize the models and the number of trees was set to 500 for all models.

Additionally, *in silico* mock communities were generated by aggregating data from axenic cultures. *In silico* mock communities were constructed by subsampling the pooled axenic data to the expected number of cells for each bacterial strain in the *in vitro* mock communities and aggregating the subsampled data for the different bacterial strains. The process of subsampling was repeated 10 times for each mock community and the repeats were pooled for making predictions.

Ultimately, the community composition of *in vitro* mock communities, *in silico* mock communities and co-cultures was then predicted using the ‘*RandomF_predict*’ function of the Phenoflow package (version 1.1.2).

#### Processing of amplicon reads

Amplicon sequences were analyzed using the DADA2 package (version 1.30.0) in R as described by Callahan et al. (62). After filtering, the average number of reads per sample was 48368.

#### Comparison of abundances

The performance of the techniques in determining the relative community composition was assessed by the root mean squared error (RMSE):

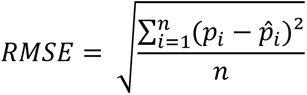

where *p* represents the known relative abundance, *p^* the predicted relative abundance, and *n* the total number of bacterial species. When a prediction is close to the actual abundance, the metric will be close to zero. Since this metric can only be calculated when the actual relative abundances are known, the calculations were only performed for the mock communities.

## Results

### Performance of random forest classifiers trained on flow cytometry data

To be able to determine the community composition of synthetic communities through flow cytometry, random forest classifiers were trained on flow cytometry data from axenic cultures (Supplementary 4). To evaluate how performance of random forest classifiers was affected by the number of species and the phylogenetic distance between the species in the classifiers, the accuracy of the classifiers was determined on 25% of the data (testing set) (Figure 3). In general, the more bacterial strains included in the model, the lower the accuracy. At the same time the relative increase compared to random guessing was higher when there were more strains included in the model. The model which included 7 bacterial strains, all oral streptococci, showed lower accuracy compared to the models which included 6 or 8 bacterial strains. The community members of the model with 7 strains also showed less phylogenetic difference (*Mock 5*, Figure 2).

**Figure 3.**
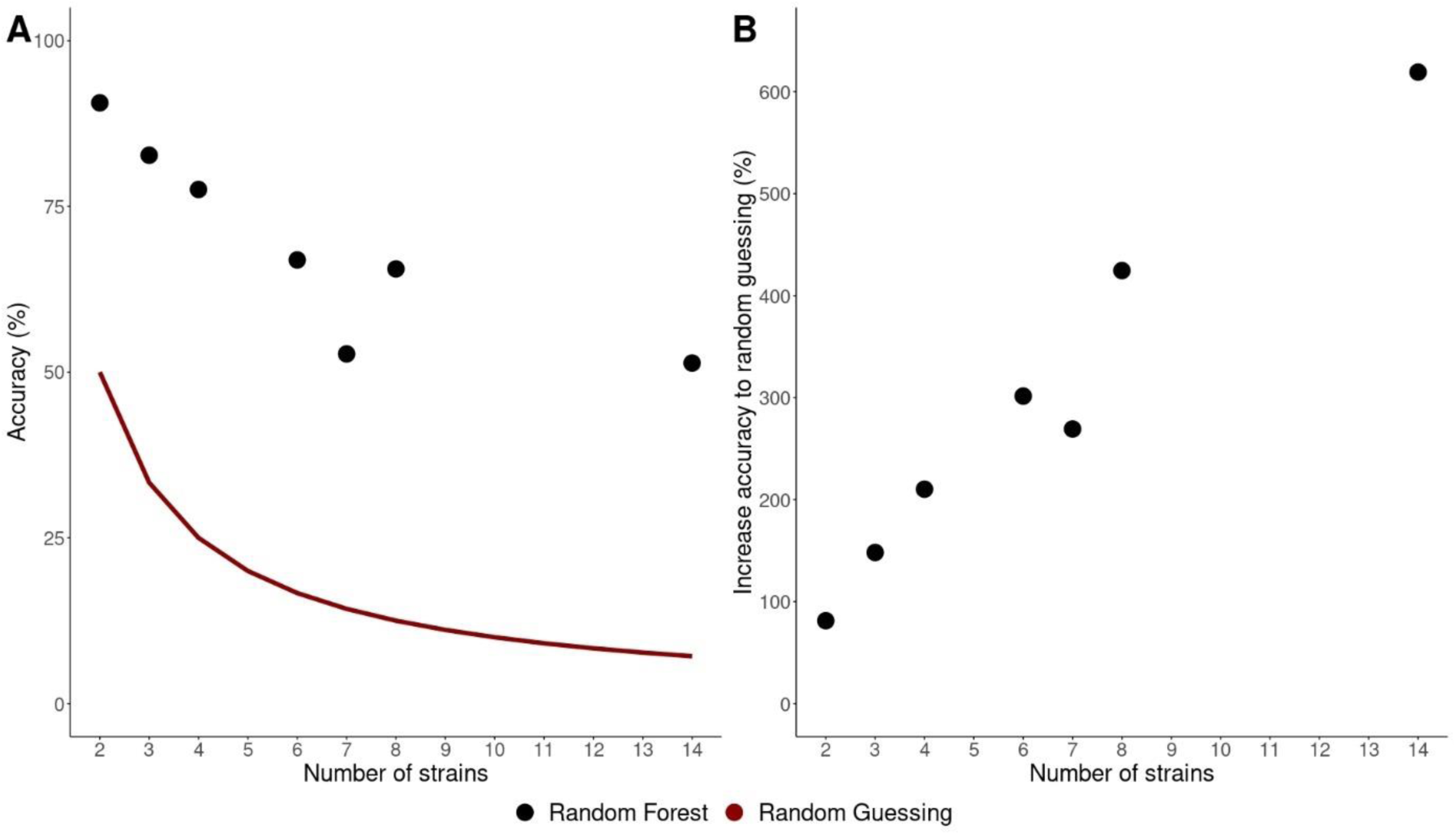
Accuracy of random forest models trained on axenic cultures. The strains used in the different models are the strains used for preparing the mock communities (Figure 2; Supplementary 4 Table S3). (A) Accuracy as a function of the number of strains in the model. The red line indicates the accuracy in case of random guessing. (B) The increase in accuracy relative to random guessing for each model.

Confusion matrices are listed in Supplementary 4. The classifiers trained on *S. oralis*, *F. nucleatum*, *P. gingivalis* and/or *V. parvula* had evenly distributed class accuracies for the different bacterial strains, with a maximum difference in class accuracy of 5.8% (Supplementary 4 Figure S3; Supplementary 4 Figure S4; Supplementary 4 Figure S5). However, for the other classifiers, class accuracy is more dependent on the bacterial strain. *P. intermedia* is predicted with high accuracy throughout the different classifiers (> 82%). For the oral streptococci, *S. salivarius* (class accuracy 71%) and *S. gordonii* (class accuracy 71%) were the easiest to differentiate, while *S. sobrinus* (class accuracy 33%) was the hardest to differentiate (Supplementary 4 Figure S7). Additionally, *S. mutans* and *S. sobrinus* were often confused with each other (Supplementary 4 Figure S7; Supplementary 4 Figure S8). *P. gingivalis* and *A. actinomycetemcomitans* were also sometimes confused with each other (Supplementary 4 Figure S8; Supplementary 4 Figure S9). Remarkably, *A. naeslundii* and *A. viscosus* were relatively easy to distinguish from each other, given that they belong to the same genus (Supplementary 4 Figure S6; Supplementary 4 Figure S9).

### Quantification of mock communities

To establish how flow cytometry compared to PCR-based methods, the community composition of mock communities was determined through flow cytometry, strain specific qPCR and 16S rRNA gene amplicon sequencing. First, the performance of flow cytometry on *in silico* and *in vitro* mock communities was evaluated, so that the effect of preparing the mock communities *in vitro* could be assessed. *In silico* mock communities were prepared in the same concentrations as the *in vitro* mock communities by combining flow cytometry data from axenic cultures. The predictions made for the *in silico* mock communities closely resembled the theoretically calculated situation (Figure 4). The theoretically calculated abundances were based on the flow cytometric measurements of the axenic cultures (Supplementary 1). The RMSE of the predictions for the *in silico* mock communities were lower compared to the RMSE for the *in vitro* mock communities, thus the performance was better for *in silico* mock communities. However, when greater numbers of strains were included in the mock communities, the RMSE of the predictions for the *in silico* and *in vitro* situation became more alike. Moreover, if an uneven mock community was considered (*Mock 8* and *Mock 9*), the predictions from the random forest classifiers yielded bigger errors. Analysis of the relative abundance of singlets (i.e. an event measured by flow cytometry that is a single cell) in the different mock communities revealed that the *in silico* situation coincides with the theoretically calculated relative abundance of singlets. However, this was not the case for the *in vitro* situation, where the relative abundance of singlets was higher compared to the theoretically calculated value, except for *Mock 2* (Supplementary 5).

**Figure 4.**
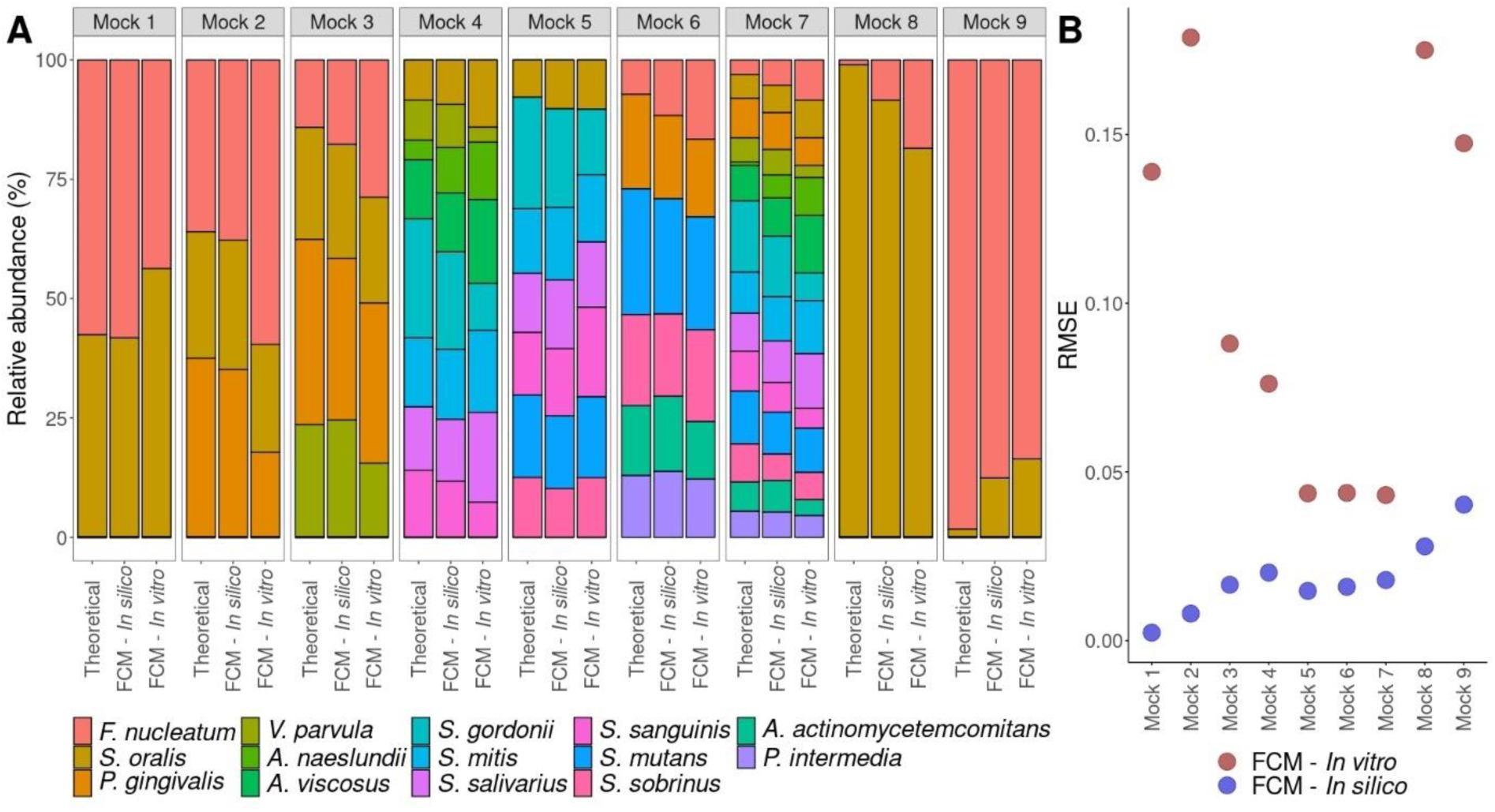
Mock communities as predicted by flow cytometry. (A) The relative abundance of the bacterial strains for different situations. ‘Theoretical’ refers to the theoretically calculated relative abundance, ‘FCM – *In silico*’ to the prediction of an *in silico* constructed mock community prepared by combining flow cytometry measurements of axenic cultures, and ‘FCM – *In vitro*’ to the prediction based on a flow cytometric measurement of an *in vitro* prepared mock community. (B) The RMSE of the prediction of the relative abundance compared to the theoretically calculated relative abundance for both the *in vitro* (red) and *in silico* (blue) mock communities.

Second, *in vitro* mock communities were quantified through flow cytometry, strain specific qPCR, and 16S rRNA gene amplicon sequencing (Figure *5*). For all mock communities, the flow cytometry model including only the strains present in the mock community in question performed best (lowest RMSE) compared to the flow cytometry models that had additional strains or strains missing to the mock community. When a flow cytometry model included more strains than those present in the mock community, the model wrongly annotated part of the community as strains which were not present in the mock community (7.7 – 21.0%). In case the model was trained on fewer strains than the ones present in the mock community, the model failed to identify the missing strains. For the even mock communities (*Mock 1*, *Mock 2* and *Mock 3*), the predictions based on flow cytometry (model with correct strains) resembled the theoretically calculated relative abundances best. Furthermore, the RMSEs of the flow cytometry models trained on more strains than the strains present in the mock community were lower or similar compared to the RMSEs of the PCR-based methods. When an uneven mock community was considered (*Mock 8* and *Mock 9*), strain specific qPCR was closest to the theoretical calculated relative abundances. Additionally, the RMSE of 16S rRNA gene amplicon sequencing was most consistent over the different mock communities. Last, *S. oralis* was overinflated in all mock communities when using qPCR.

**Figure 5.**
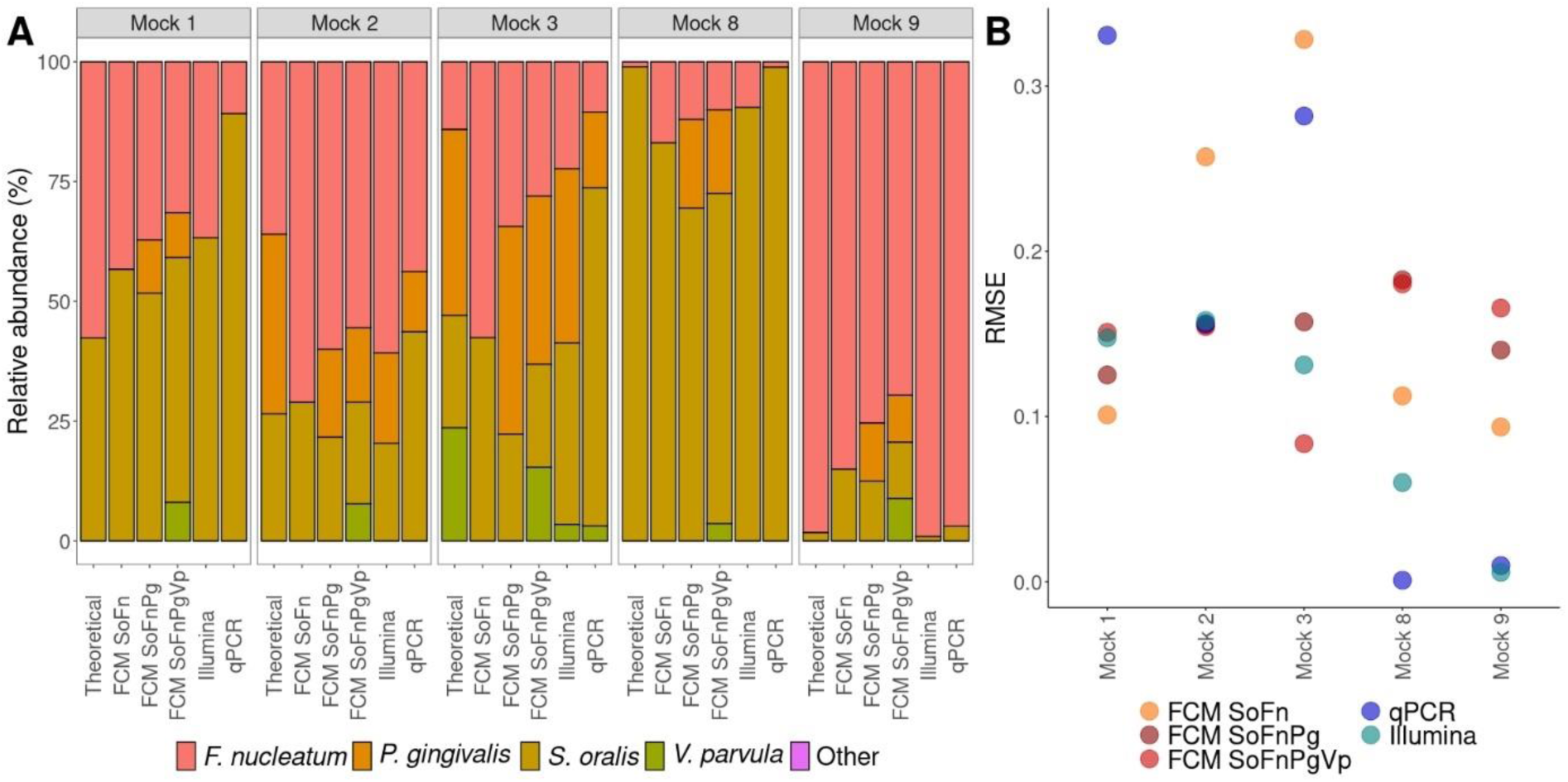
Relative quantification of mock communities determined through different techniques. ‘FCM’ refers to predictions made based on flow cytometric measurements of the mock communities. ‘SoFn’ refers to the random forest model trained on *S. oralis* and *F. nucleatum*, ‘SoFnPg’ to the model trained on *S. oralis*, *F. nucleatum* and *P. gingivalis*, ‘SoFnPgVp’ to the model trained on *S. oralis*, *F. nucleatum*, *P. gingivalis* and *V. parvula*. (A) Relative abundance of the different mock communities determined through flow cytometry (FCM), 16S rRNA gene amplicon sequencing (Illumina) and strain specific qPCR. ‘Theoretical’ indicates the theoretically calculated relative abundance. (B) RMSE of each technique compared to the theoretically calculated relative abundance.

When absolute abundances were examined, qPCR led to cell concentrations of about 10-fold compared both to cell concentrations obtained from flow cytometry and the theoretically calculated cell concentrations (Figure 6A). The theoretically calculated cell concentrations were based on flow cytometric measurements of the axenic cultures. Cell concentrations determined through qPCR were obtained by considering the sample volume taken for DNA extraction and the gene copy number for the targeted gene of each bacterial strain. Comparison of cell concentrations determined through flow cytometry and the theoretical calculation showed that flow cytometry led to lower cell concentrations, except in *Mock 2* (Figure 6B). To assess whether part of the difference in cell concentrations could be explained by the occurrence of cell aggregates, singlet analysis was performed. It revealed that about 70% of measured events through flow cytometry were singlets (Supplementary 5).

**Figure 6.**
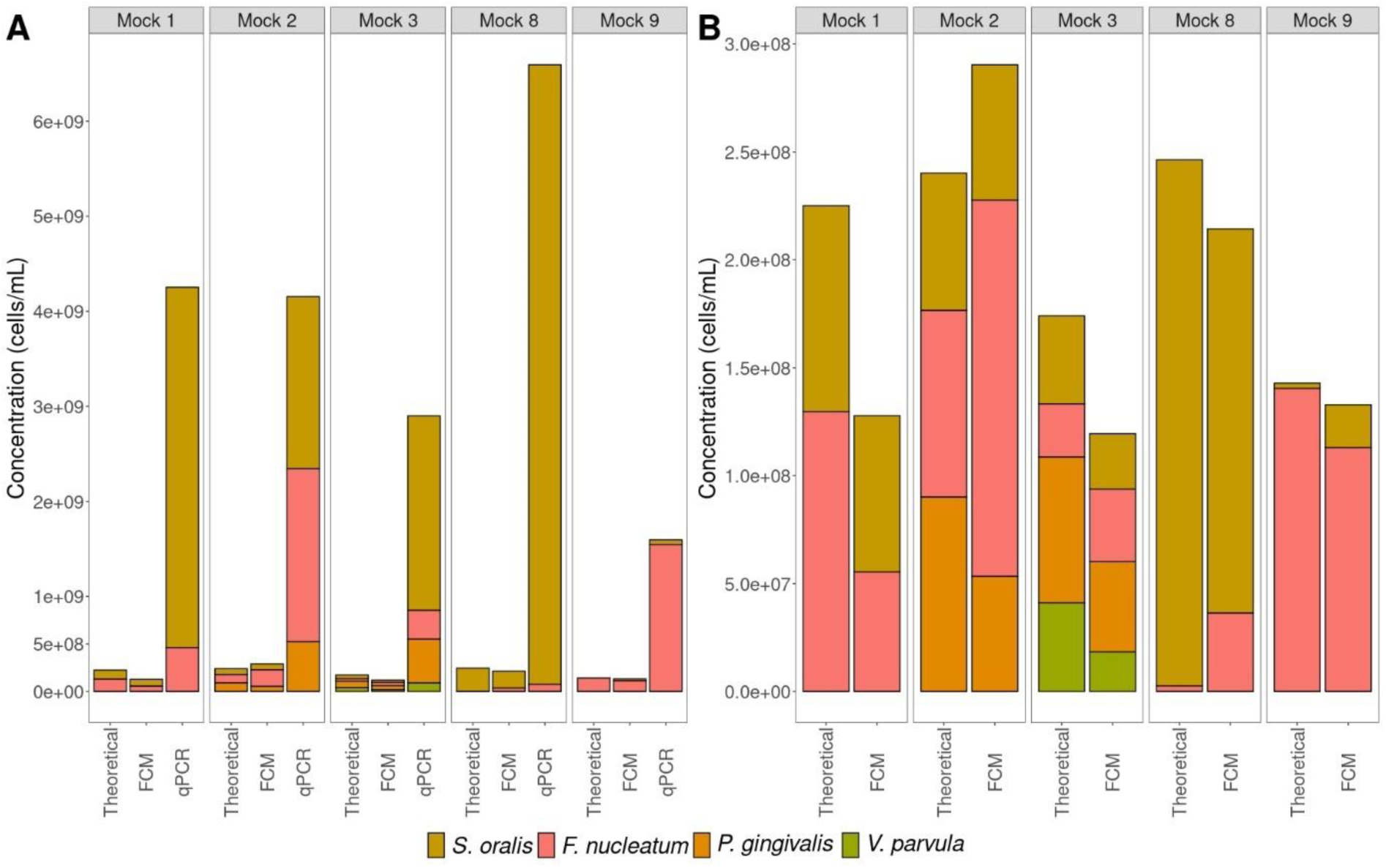
Absolute quantification of mock communities through flow cytometry and qPCR. ‘Theoretical’ indicates the theoretically calculated absolute abundance and ‘FCM’ the absolute abundance determined through flow cytometry. (A) Flow cytometry and strain specific qPCR compared to the theoretically calculated situation. (B) A more detailed depiction of the results of flow cytometry compared to the theoretically calculated situation.

### Quantification of co-cultures

Since phenotypes of bacteria can change when grown together, the community composition of co-cultures was quantified using flow cytometry, strain specific qPCR, and 16S rRNA gene amplicon sequencing. The co-cultures differed from the mock communities as these were grown together, where the mock communities were mixtures prepared post-growth. The relative abundances of biological replicates were similar to each other and changes in community structure between 24h and 48h were observed irrespective of the technique used (Figure 7). However, the different techniques resulted in different community compositions.

**Figure 7.**
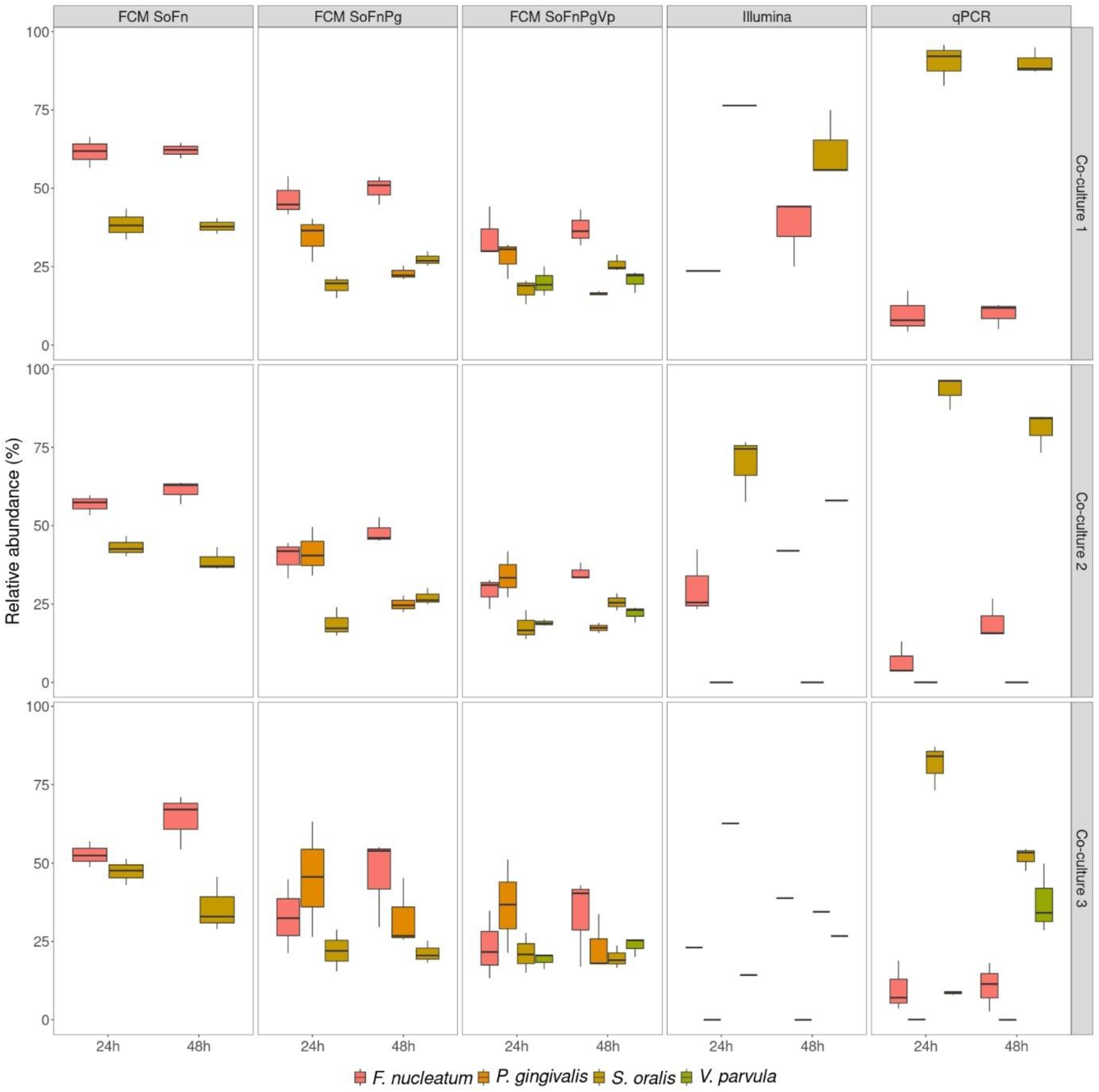
Relative abundances of co-cultures as determined through different techniques after 24h and 48h of growth. *Co-culture 1* contained *F. nucleatum* and *S. oralis*, *Co-culture 2* contained *F. nucleatum*, *P. gingivalis* and *S. oralis*, and *Co-culture 3* contained *F. nucleatum*, *P. gingivalis*, *S. oralis* and *V. parvula*. ‘FCM’ refers to predictions made based on flow cytometric measurements of the co-cultures. ‘SoFn’ refers to the random forest model trained on *S. oralis* and *F. nucleatum*, ‘SoFnPg’ to the model trained on *S. oralis*, *F. nucleatum* and *P. gingivalis*, ‘SoFnPgVp’ to the model trained on *S. oralis*, *F. nucleatum*, *P. gingivalis* and *V. parvula*. With 16S rRNA gene amplicon sequencing (Illumina), only one replicate was sequenced for *Co-culture 1* after 24h of growth, *Co-culture 2* after 48h of growth and *Co-culture 3* for both timepoints.

When using flow cytometry to determine the community composition, models trained on more strains than the ones present in the co-culture wrongly annotated part of the community as strains that were not present (19.0 – 47.9%). When comparing the co-cultures to the mock communities consisting of the same bacterial strains (Figure 5A; Figure 7), the relative abundance of incorrectly predicted strains through flow cytometry was higher with the co-cultures. For example, *Mock 1*, *Mock 8*, *Mock 9*, and *Co-culture 1* only contained *S. oralis* and *F. nucleatum*. However, when the flow cytometry model that also includes *P. gingivalis* and *V. parvula* was used to predict the relative abundance of the strains, a higher combined relative abundance of *P. gingivalis* and *V. parvula* was found in the co-cultures (*Mock 1*: 17.4%, *Mock 8*: 21.0 %, *Mock 9*: 18.6%, *Co-culture 1 24h*: 47.9%, *Co-culture 1 48h*: 37.1%).

Analysis of absolute abundances for the co-cultures revealed that qPCR yielded roughly 10-fold higher cell concentrations compared to flow cytometry (Figure 8), except for *Co-culture 2A* after 48h of growth. According to qPCR, *Co-culture 1* and *Co-culture 3*, except for replicate A, showed an increase in total cell concentration between 24h and 48h. For *Co-culture 2* a decrease was observed. When statistical differences in total cell concentrations between timepoints were assessed, there was a significant difference in cell concentration only for *Co-culture 1* between timepoints (Supplementary 6). According to flow cytometry, total cell concentrations between timepoints remained stable for *Co-culture 1* and *Co-culture 2*, while a decrease was observed for *Co-culture 3*. The decrease was not found to be statistically significant in a two-sided paired t-test (Supplementary 6). Last, an increase in relative abundance of singlets between timepoints was observed for all replicates of all co-cultures, except for *Co-culture 1A* and *Co-culture 2A* (Supplementary 5).

**Figure 8.**
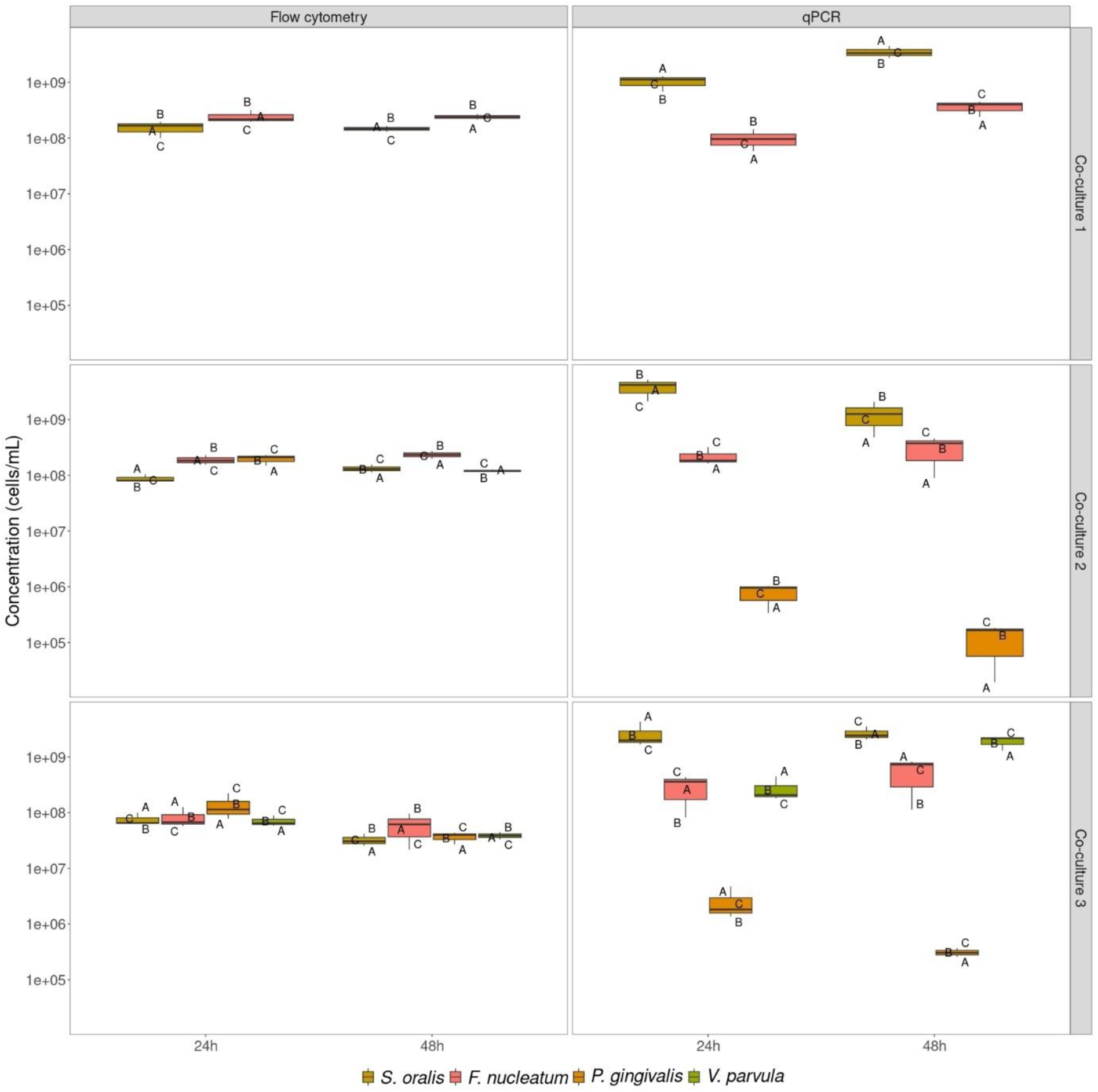
Absolute abundances of co-cultures after 24h and 48h of growth as determined by flow cytometry and strain specific qPCR. For flow cytometry, only the random forest model containing the inoculated strains for each co-culture was used to determine cell concentrations. The letters ‘A’, ‘B’ and ‘C’ show the concentrations for each biological replicate.

## Discussion

Quantification of community structure of synthetic communities is necessary to allow for descriptive and mechanistic research of microbial consortia (15, 16, 63). Quantification typically is performed through labor-intensive and/or expensive techniques such as plate counting, strain specific qPCR and 16S rRNA gene amplicon sequencing. In this study we investigated if flow cytometry can be used as a fast and cheap alternative to quantify the composition of synthetic mock communities and co-cultures and compared its performance against strain specific qPCR and 16S rRNA gene amplicon sequencing. Our results show that flow cytometry outperformed the PCR-based methods when it came to determining the community composition of *in vitro* mock communities, but that the performance decreased when it was used to determine the composition of bacteria that had been grown in co-culture.

The use of flow cytometry to quantify synthetic communities has been demonstrated for *in silico* mock communities before (37). In line with these findings, we found that the more bacterial strains included in the flow cytometry based random forest classifiers, the lower the accuracy of the classifiers (Figure 3A). The classifier that included only seven oral streptococci yielded lower classifier performance than the classifier that included eight bacterial strains of varying genera. This can be attributed to the phylogenetic related species having similar phenotypes. Thus, it is more difficult to differentiate them (64). Since previous research showed that shape and size differences were not enough to fully explain the individual classification accuracies of bacterial strains and that differences in genome size between different bacterial strains change the phenotypic fingerprint, more research is needed to clarify why certain strains are easier to differentiate (38, 55). Simultaneously, the relative proportion of the accuracy of the models compared to random guessing increased when more strains were included (Figure 3B). Therefore, the predictions may be better when more strains are included, at least up to a total number of strains of 14 as shown by our results.

We found that the flow cytometry-based predictions for *in silico* prepared mock communities were better than the predictions made for the same mock communities prepared *in vitro*, which can be rationalized by both technical and biological variation (65–67). Yet, the difference between the RMSE for the *in* silico and *in vitro* mock communities became smaller when more strains were included (Figure 4). In general, analysis of singlets showed a higher abundance of singlets in the *in vitro* mock communities (Supplementary 5). Singlets can be considered single cells, while multiplets (non-singlets) are aggregates of cells. Our experimental procedure for creating mock communities included additional vortexing compared to the axenic cultures, which likely led to a reduction in aggregates. However, for *Mock 2*, which consisted of *S. oralis*, *F. nucleatum* and *P. gingivalis*, this was not observed. Co-aggregation between *F. nucleatum* and *P. gingivalis* may explain this observation and is well described in literature (68–71). Moreover, we found that in case of an uneven mock community, the predictions were less accurate for both the *in silico* and *in vitro* condition and that this is strain dependent (Figure 4). Thus, we suggest taking caution when predicting the composition of uneven mock communities using flow cytometry.

When comparing flow cytometry to PCR-based methods for quantifying mock communities, we observed that flow cytometry performed better (i.e. lower RMSE) than both strain specific qPCR and 16S rRNA gene amplicon sequencing for even communities (Figure 5). Better performance of flow cytometry compared to 16S rRNA gene amplicon sequencings was reported for dual-strain mock communities of gut bacteria (55). Previous reports indicated that the result of the determination of the microbial community structure is dependent on DNA extraction method, primer choice and sequencing platform, and that 16S rRNA gene amplicon sequencing does not always correspond well to the actual bacterial communities (72–75). This could explain why flow cytometry performed better. As was expected, the flow cytometry-based classifier that only included the relevant strains for the mock community resulted in the best prediction. In addition, performance of the flow cytometry-based classifiers that were trained on more strains than the strains present in the even mock communities was better compared to the PCR-based methods. Thus, it can be relevant to consider a model that includes more strains than the strains present in a synthetic community. For example, when limited computational power is available, or when trying to achieve fast screening of community composition and time constraints do not allow examination of each community through different models. Overall, the RMSE of 16S rRNA gene amplicon sequencing was lower compared to strain specific qPCR and thus seemed more accurate for the determination of the community composition. Still, it must be noted that the relative abundance of *S. oralis* was inflated for all mock communities when using strain specific qPCR compared to 16S rRNA gene amplicon sequencing. Since the target gene for strain specific qPCR and 16S rRNA gene amplicon sequencing was different for *S. oralis* between the techniques, it may be that strain specific qPCR performs better when using the same target. This is portrayed by the better RMSE for strain specific qPCR in *Mock 8*, where the relative abundance of *F. nucleatum* becomes less relevant.

Absolute quantification of mock communities revealed large differences in total cell concentrations between flow cytometry and strain specific qPCR, where qPCR leads to concentrations about 10-fold higher compared to flow cytometry (Figure 6). Part of the difference can be explained by underestimation of the actual cell count by flow cytometry due to high presence of cells aggregates that are erroneously counted as a single cell (Supplementary 5). Nonetheless, cell aggregation should not affect cell count by qPCR as quantification is based on the number of gene copies detected in the DNA extract of the sample. Accordingly, we strongly advise breaking aggregates before using flow cytometry to quantify synthetic communities. This can be done both mechanically (e.g. vortexing and sonication) and chemically (e.g. use of TWEEN), or by a combination of both (76, 77). Moreover, some of the bacteria used in the experiment are reported not to have reached stationary phase yet after 24h of growth (78, 79). Bacterial cells that are still in the exponential phase are known to have additional copy numbers of genes because they are actively dividing (80–82). This will also lead to higher absolute abundances determined through strain specific qPCR. To elucidate the discrepancies between flow cytometry and (strain specific) qPCR, additional experiments are required. Interestingly, a discrepancy was observed between the theoretical cell concentration and the cell concentration determined through flow cytometry as well (Figure 6B). This can be explained by the difference in relative abundance of singlets between the theoretical cell concentration and cell concentration determined through flow cytometry (Supplementary 5).

When co-culturing bacteria, we found that neither flow cytometry, strain specific qPCR and 16S rRNA gene amplicon sequencing resulted in similar community composition. Since our experiments with *in vitro* mock communities did not establish one of the techniques as highly accurate over the different tested conditions, none of the techniques can be used as a benchmark for the community composition of co-cultures. As was the case for the mock communities, singlet analysis revealed that a significant proportion of the microbial communities were multiplets and thus introduced bias in the community structure determined through flow cytometry (Supplementary 5). In addition, increased relative abundances of wrongly predicted strains compared to the mock communities were found when a random forest classifier was used that also included strains not in the synthetic community (mock communities: 7.7 – 21.0%; co-cultures: 19.0 – 47.9%). The wrongly predicted strains can be interpreted as additional phenotypes of the strains that make up the co-culture, which is in accordance with previous research. It was shown that phenotypes of bacteria change when co-cultured (52, 53). Therefore, we argue that because bacterial phenotypes change in co-cultures, the current flow cytometric approach is not suitable for accurate determination of the community composition in co-cultures. Likewise, literature indicates only similar trends between flow cytometry and 16S rRNA gene amplicon sequencing for co-cultures of gut bacteria (55). One solution to the issue with the flow cytometric approach could be the use of flowFISH, where fluorescent DNA probes are used to target specific RNA sequences of the different taxonomic strains (83–86). Notwithstanding, this would make the method more expensive and labor-intensive, as opposed to the approach described in this research. Alternatively, the use of specific targeted antibody labeling of bacteria, as was shown for *F. prausnitzii*, could lead to better results (87). Another solution could lie in the use of FACS. Heyse et al. showed that the presence and abundance of bacterial taxa in environmental samples could be predicted through flow cytometry. They sorted different regions in the flow cytometry data space and assessed which bacterial taxa were abundant in each region (88). A similar approach could be used here as well, so that the different regions could be assigned to the different bacterial strains making up the synthetic community. Last, similar observations as with the mock communities were made when it came to total cell concentrations (Figure 8). Again, the relative abundance of singlets could explain part of the difference between strain specific qPCR and flow cytometry (Supplementary 5). After 48h co-cultures were expected to have reached stationary phase, based on the growth curves of the individual strains. Indeed, we observed a higher relative abundance of singlets after 48h, which implies less cells actively dividing. Notably, flow cytometry and strain specific qPCR did not agree on the evolution of cell concentrations between timepoints. More investigation is needed to shed light on the reason for this difference.

To conclude, we found that flow cytometry performed better than strain specific qPCR and 16S rRNA gene amplicon sequencing when determining the community composition of mock communities, but that for co-cultures performance was reduced. The flow cytometry method was affected by the occurrence of cell aggregates, the inclusion of additional bacterial strains in the model compared to the strains present in the synthetic community and changing phenotypes of the strains when co-cultured. Considering these aspects, further development of the flow cytometry approach will increase the accuracy of the method and could show potential as a fast and cheap alternative for quantifying co-cultures. Finally, we found large discrepancies between flow cytometry and strain specific qPCR, which should be investigated further.

## Code and data availability

Raw flow cytometry data and metadata are available at FlowRepository under ID FR-FCM-Z8YN. Primer clipped sequence reads are available at NCBI as Sequence Read Archive (SRA) under BioProject ID PRJNA1112420 (http://www.ncbi.nlm.nih.gov/bioproject/1112420). The data analysis scripts are available at https://github.com/famermans/QuantificationSynCom.

## Acknowledgements

The authors would like to thank Frederiek-Maarten Kerckhof, Peter Rubbens, and Ruben Probs for their advice during data analysis, Tim Lacoere for his advice during the experimental work and for providing the figure of the experimental workflow, and Valérie Mattelin and Fien Waegenaar for critically reading the manuscript. This work was supported by Research Foundation – Flanders (FWO, Belgium) (grant number: G0B2719N).

## Author contributions

FM, IC and NB conceived and designed the study. FM and IC performed the experimental work and analyzed the data. FM and IC wrote the manuscript. FM created the figures. WT and NB supervised the findings of this work. All authors reviewed and approved the manuscript.

